# Development of alcoholic liver disease model for drug evaluation from human induced pluripotent stem cell derived liver organoids

**DOI:** 10.1101/2024.03.10.584291

**Authors:** Zhiwei Feng, Bingrui Zhou, Qizhi Shuai, Yunliang Wei, Ning Jin, Xiaoling Wang, Hong Zhao, Zhizhen Liu, Jun Xu, Jianbing Mu, Jun Xie

**Affiliations:** Department of Biochemistry and Molecular Biology, Shanxi Key Laboratory of Birth Defect and Cell Regeneration, MOE Key Laboratory of Coal Environmental Pathogenicity and Prevention, Shanxi Medical University, Taiyuan 030001, Shanxi, China; Department of Hepatobiliary and Pancreatic Surgery and Liver Transplant Center, the First Hospital of Shanxi Medical University, Taiyuan 030001, China; Laboratory of Malaria and Vector Research, National Institute of Allergy and Infectious Diseases, National Institutes of Health, 12735 Twinbrook Parkway, United States

**Keywords:** Human induced pluripotent stem cells, Liver organoids, Alcoholic liver disease, Drug screening

## Abstract

Alcoholic liver disease (ALD) poses a significant health challenge, demanding comprehensive research efforts to enhance our comprehension and treatment strategies. However, the development of effective treatments is hindered by the limitations of existing liver disease models. Liver organoids, characterized by their cellular complexity and three-dimensional (3D) tissue structure closely resembling the human liver, hold promise as ideal models for liver disease research. In this study, we employ a meticulously designed protocol involving the differentiation of human induced pluripotent stem cells (hiPSCs) into liver organoids. This process incorporates a precise combination of cytokines and small molecule compounds within a 3D culture system to guide the differentiation process. Subsequently, these differentiated liver organoids are subjected to ethanol treatment to induce ALD, thus establishing a disease model. Rigorous assessment through a series of experiments reveals that this model partially replicates key pathological features observed in clinical ALD, including cellular mitochondrial damage, elevated cellular reactive oxygen species (ROS) levels, fatty liver, and hepatocyte necrosis. In addition, this model offers potential utility in screening drugs for ALD treatment. Taken together, the liver organoids model of ALD, derived from hiPSCs differentiation, emerges as an invaluable platform for advancing our understanding and management of ALD in clinical settings.

## Introduction

The report from the World Health Organization indicates that alcohol abuse led to over 3 million deaths in 2018, accounting for 1 in 20 deaths and contributing to more than 5% of the global disease burden. Prolonged alcohol abuse can lead to the development of Alcoholic Liver Disease (ALD), which encompasses fatty liver (steatosis), alcoholic hepatitis, and alcoholic cirrhosis[1, 2]. The pathological features of ALD include hepatocyte steatosis, oxidative stress, inflammation, and liver fibrosis. Progression of the disease can lead to hepatocyte necrosis and apoptosis, liver cirrhosis, and even liver cancer [3, 4]. At present, the models used to study ALD mainly include animal models and liver cell lines culture models [5, 6]. Two-dimensional cultures of liver cell lines have traditionally been used to mimic diseases and develop medications. For instance, flat cultures of liver stellate cells (HSCs) have been employed, with or without the addition of soft hydrogels such as polyacrylamide gels. However, the genetic profiles of these cells often do not align with those of cirrhotic tissues in humans. Translating findings from 2D cultures has been challenging due to their inability to replicate natural characteristics such as dynamic physical and chemical signals and the microenvironmental structures present in the liver lobule, which often leads to a rapid decline in liver cell function[7]. Therefore, the application of these models in ALD research is limited due to species differences and the constraints of two-dimensional (2D) culture [8–10].

Organoids refer to the self-assembly of adult stem cells or pluripotent stem cells in a 3D *in vitro* culture environment to form tissue analogs with 3D structures that accurately reflect the characteristics of the original tissue (derived from adult stem cells) or directed differentiated tissue (derived from induced pluripotent stem cells) [11]. Organoids originating from iPSCs are widely recognized as a pivotal element in the field of disease modeling based on organoids. Recent studies have demonstrated the successful development of vascularized organoids derived from iPSCs. In one specific study, integration of stromal elements such as vasculature, fibroblasts, and immune cells was achieved by utilizing mesodermal progenitor cells induced from iPSCs [12]. Organoid culture offers the potential for long-term culture *in vitro* expansion while maintaining stable genetic characteristics, thus providing a solution to the challenges of liver disease models [13]. *In vitro* organoid models represent a significant advancement for translational research, as these 3D models closely mimic *in vivo* biological processes such as tissue renewal and the response of tissues to drugs, toxins, and mutagenesis. Compared to traditional monolayer cultures, 3D liver organoid cultures offer more accurate models for studying hepatic toxicity.

In this study, we initially differentiated hiPSCs into liver organoids. These derived liver organoids underwent assessment for their structural integrity, cellular composition, expression of liver-related proteins, and specific liver functions. Subsequently, the liver organoids were subjected to treatment with different concentrations of ethanol to establish an ALD model. This model successfully replicated key pathological features of ALD, including hepatocyte adiposeness, mitochondrial damage, elevated cellular ROS, and cell death. Finally, we selected three drugs with reported therapeutic and preventive potential for ALD, utilizing the established liver organoids model as a platform for drug screening and evaluating its applicability for the clinical ALD study.

## Materials and Methods

### Cell lines and Cell culture

The human induced pluripotent stem cell line, hiPSC-B1, was purchased from Cellapy^®^ (CA4025106, Beijing, China) and cultured on plates coated with low growth factor Matrigel (354230, Corning, USA) using mTeSR^TM^ 1 medium (85850, StemCell, Vancouver, Canada) supplemented with 100 U/ml penicillin/streptomycin (Solarbio, Beijing, China). The culture was maintained in a humidified atmosphere containing 5% CO_2_ at 37 L in a cell incubator (Eppendorf, Germany). The medium was changed daily, and the cells were passaged at a ratio of 1:5-1:8 every 4-7 days.

### Liver organoids Generation

Undifferentiated hiPSC-B1 cell lines with passage numbers between 30–50 were digested into single cells by Accutase (00-4555, Thermo Fisher, USA) and seeded onto Matrigel-coated culture plates. Once the cells reached approximately 80% confluent without signs of differentiation, the differentiation process was initiated. The medium was replaced with RPMI 1640 medium (31800, Solarbio, Beijing, China) containing 100 ng/mL Activin A (120-14E, PeproTech, USA) and 50 ng/mL bone morphogenetic protein 4 (BMP4; 314-BP, R&D Systems, USA) on day 1, followed by RPMI 1640 medium containing 100 ng/mL Activin A and 0.2% Knockout serum replacement (KSR; A3181502, Gibco, USA) on day 2, and RPMI 1640 medium containing 100 ng/mL Activin A and 2% KSR on day 3. From day 4 to day 6, cells were cultured in High Glucose DMEM medium (SC102-02, Sevenbio, Beijing, China) supplemented with 50 ng/mL recombinant human fibroblast growth factor 10 (FGF-10; 100-26, PeproTech, USA), 3 μM CHR99021 (SML1046, Sigma, USA), 10% KSR, 1% NEAA (11140, Gibco, USA). The cells were maintained at 37 L in 5% CO_2_ with 95% air in the cell incubator, with daily medium changes. By approximately day 6, the spheroids of foregut endoderm appeared on the plate. On day 7, the spheroids were dissociated with Accutase into single cells and embedded in 100% Matrigel drops on the plates in High Glucose DMEM medium containing 3 μM CHR99021, 5 ng/mL FGF-2 (100-18B, PeproTech, USA), 0.5 μM of A83-01 (HY10432, MCE, USA) and 20 ng/mL of recombinant human epidermal growth factor (EGF; AF100-15, PeproTech, USA), 10% KSR and 1% NEAA, and cultured for 2 days. On day 9, the medium was switched to High Glucose DMEM medium containing 2 μM retinoic acid (RA; R2625, Sigma, USA), 10% KSR, and 1% NEAA, and cultured for 4 days, with medium renewal on day 11. After RA treatment, the liver organoids were gradually formed. On day 13, the medium was switched to Hepatocyte Culture Medium (HCM; CC-3198, Lonza, Switzerland) with 10 ng/mL hepatocyte growth factor (HGF; 100-39, PeproTech, USA), 0.1 mM Dexamethasone (Dex; D4902, Sigma, USA) and 20 ng/mL Oncostatin M (OSM; 300-10, PeproTech, USA). The liver organoids were isolated from Matrigel by scraping and pipetting and passaged on day 16, with medium changes every 3 days. The liver organoids were ready for downstream applications after day 22-25.

### RNA Isolation, Reverse transcription PCR, Quantitative PCR

Total RNA was extracted from liver organoids using RNAiso Plus reagent (TaKaRa, Japan) in combination with chloroform and isopropanol. Complementary DNA (cDNA) was synthesized using the PrimeScript™ RT reagent Kit with gDNA Eraser Kit (TaKaRa, Japan) following the manufacturer’s protocol. Quantitative PCR (qPCR) was performed on an Applied Biosystems QuantStudio 3 Real-Time PCR System (Thermo Fisher, USA) using SYBR^®^ Premix Ex Taq^TM^ II (TaKaRa, Japan). All primer information for each target gene was obtained from the Primerbank website(https://pga.mgh.harvard.edu/primerbank/). The PCR reaction conditions were as follows: initial denaturation at 95 °C for 1 minute, followed by 40 cycles of 95 °C for 5 s, and 60 °C for 30 s, with final Melt Curve Stage. All samples were run in triplicate concurrently. The values were normalized using internal control products of glyceraldehyde-3-phosphate dehydrogenase (GAPDH). A complete list of the primers used is provided in Table 1.

**Table 1.**
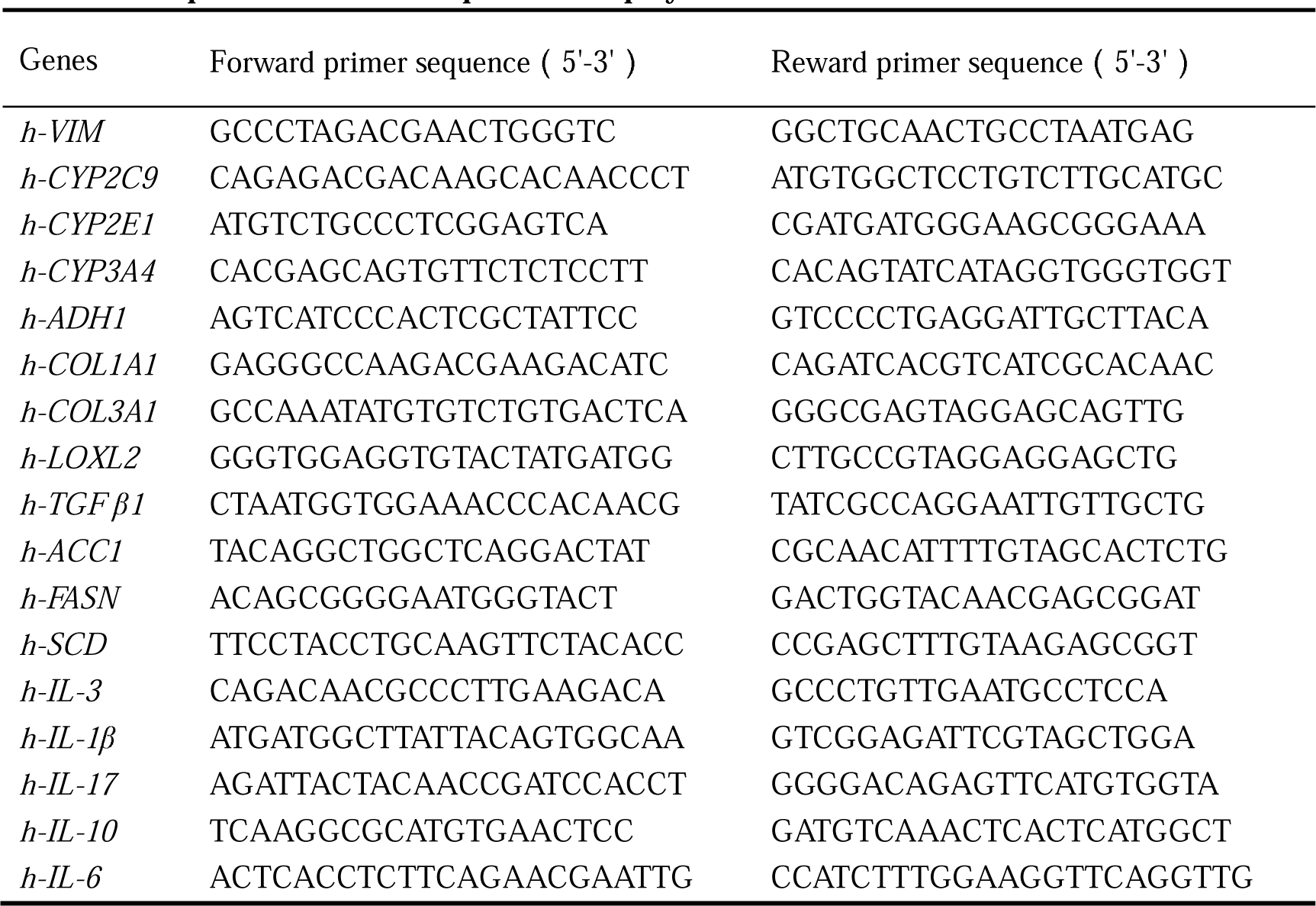
The primers used in the quantitative polymerase chain reaction.

### Immunofluorescence (IF)

The immunofluorescence staining method was conducted according to previously described protocol [14]. Liver organoids were fixed in 4% PFA for 4 hours, dehydrated, embedded in paraffin, and sectioned at 4 μm thickness. Paraffin sections were then dewaxed with xylene, rehydrated, and subjected to heat-mediated antigen retrieval by microwaving for 15 minutes in Tris-EGTA buffer (pH 9.0). Following blocking with 1% BSA/PBS (Sigma-Aldrich, USA) for 1 hour, the sections were incubated with primary antibodies in 1% BSA/PBS at 4 L overnight and secondary antibodies in 1% BSA/PBS for 1 hour in the dark at RT. Subsequently, 4’6-diamidino-2-phenylindole (DAPI; Abcam, UK) was used to counterstain the nuclei. Sections were washed 3 times with PBS for 5 minutes each, with shaking, after primary and secondary antibody and DAPI staining. Immunofluorescence images were acquired using Fluorescence Light Microscopes (Nikon, Japan). NIS-Elements Viewer software (Nikon) was used for image processing. A complete list of primary and secondary antibodies used is provided in Table 2.

**Table 2.**
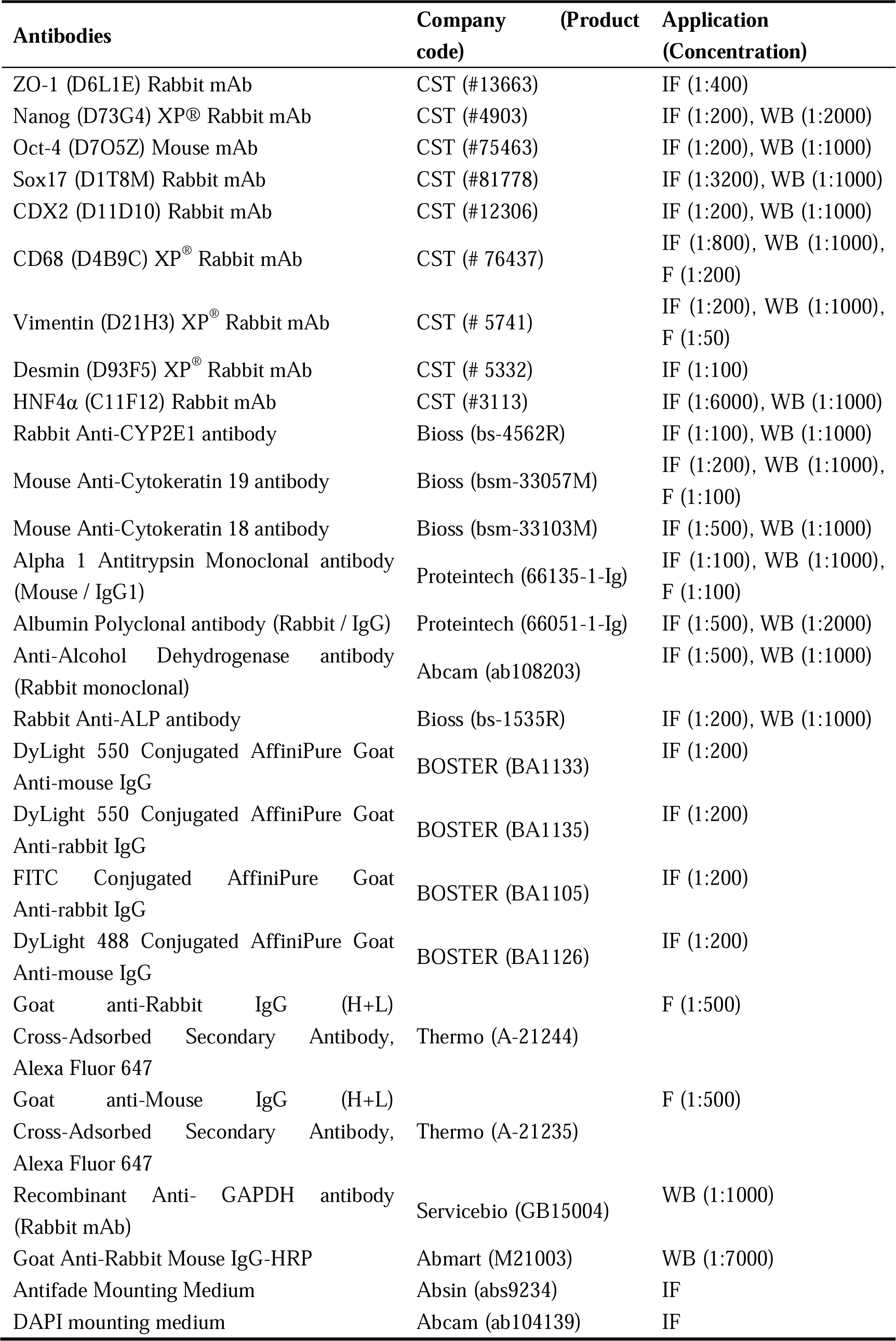
Antibodies used in immunofluorescence staining (IF), Flow cytometry (F) and Western Blotting (WB).

### Flow cytometry

Liver organoids was treated with trypsin EDTA (0.25%, Solarbio, Beijing, China) for 30 minutes and dissociated into single cells, which were then passed through 70 μm cell strainers. Subsequently, the hepatocyte-like organoid (HLO) cells were fixed in 4% PFA for 20 minutes, permeabilized in 0.2% Triton X/PBS for 10 minutes, and blocked with 1% BSA/PBS blocking solution for 30 minutes before incubating with primary antibodies for 1 hour. Following incubation with secondary antibodies for 0.5 hours in the dark, the stained cells were analyzed using BD FACS Celesta ^TM^ (BD Biosciences, USA), and the obtained data were analyzed using the FlowJo software. A complete list of primary and secondary antibodies used is provided in Table 2.

### Western blotting

Total proteins were extracted from the liver organoids using the Cell lysis buffer (Solarbio, Beijing, China) supplemented with protease inhibitors at 4 °C, and the concentration of the protein samples was quantified using the BCA Protein Assay Kit (EpiZyme, Shanghai, China). The protein samples were separated by electrophoresing in 10% sodium dodecyl sulfate-polyacrylamide gels (SDS-PAGE) and transferred onto polyvinylidene fluoride (PVDF) membranes. After blocking with 5% non-fat milk for 1 hour at room temperature, the PVDF membranes were incubated with diluted primary antibodies at 4 °C overnight. Then, the membranes were washed 3 times with Phosphate buffer saline with Tween-20 (PBST) and incubated with the peroxidase-conjugated goat anti- rabbit & mouse secondary antibody (Abmart, Shanghai, China) at room temperature for 2 hours. After three washes with PBST, the protein blots on the PVDF were detected using ECL luminous solution (EpiZyme, Shanghai, China) in ChemiDoc MP electrophoretic imaging system (Bio-Rad, USA), and the protein levels were quantified using Fiji software. A complete list of primary and secondary antibodies used is provided in Table 2.

### Indocyanine green (ICG) uptake and release assay

The liver organoids were incubated with 1 mg/ml indocyanine green (ICG; Yuanye Bio, Shanghai, China) in the medium for 30 minutes at 37 L. Following incubation, the medium containing ICG was discarded and the organoids were washed 3 times with PBS. The uptake of ICG was examined under the inverted microscope (Leica, Germany). Subsequently, the organoids were returned to the culture medium and incubated for an additional 48 hours to assess the release of cellular ICG.

### Periodic acid-Schiff staining, Sirius red staining, Masson’s Staining

Liver organoids paraffin-embedded sections were subjected to staining with Periodic acid-Schiff (PAS; Servicebio, Wuhan, China), Sirius red (G-clone, Beijing, China), and Masson’s trichrome (Servicebio, Wuhan, China) according to the manufacturer’s instructions. The collagen volume fraction (CVF) was quantified using Fiji software.

### Intracellular lipids assays

The intracellular neutral lipid contents were measured using the lipophilic fluorescent probe dipyrromethene boron difluoride (BODIPY; GLPBIO, Tianjin, China) and the nuclear dye Hoechst 33342 (Solarbio, Beijing, China). Liver organoids were dissociated into single organoids and incubated with Hepatocyte Culture Medium (HCM) containing 2 μM BODIPY and 10 μM Hoechst 33342 for 20 minutes at 37 °C in the dark. After incubation, the organoids were washed three times with PBS, resuspended in PBS with Antifade Mounting Medium (Absin), and observed using inverted fluorescence microscopy. BODIPY was visualized using the green fluorescence channel, while Hoechst 33342 was visualized using the blue fluorescence channel. The organoids suspension was then transferred to a 96-well or 384-well microtiter plate, and the fluorescence intensity was detected using a SpectraMax iD3 microplate reader (Molecular Devices, USA). The excitation wavelength and emission wavelength of Hoechst 33342 are 350 nm and 460 nm, respectively. The excitation and emission wavelengths of BODIPY are 488 nm and 530 nm, respectively. The intracellular neutral lipid accumulation is indicated by an increase in the green/blue fluorescence intensity ratio.

### Mitochondrial membrane potential (MMP) assay

Alterations in mitochondrial membrane potential were detected using JC-1 kit (Solarbio, Beijing, China) according to the manufacturer’s instructions. Liver organoids were dissociated into single organoids and incubated with JC-1 work solution for 30 minutes at 37 °C in the dark. Afterward, they were washed three times with JC-1 buffer, resuspended in PBS with Antifade Mounting Medium, and observed using inverted fluorescence microscopy. JC-1 monomer was visualized with the green fluorescence channel, while JC-1 aggregates were observed using the red fluorescence channel. The organoids suspension was then transferred to a 96-well or 384-well microtiter plate, and the fluorescence intensity was detected on the SpectraMax iD3 microplate reader. The excitation wavelength and emission wavelength of the JC-1 monomer are 488 nm and 530 nm, respectively. The excitation and emission wavelengths of JC-1 aggregates are 529 nm and

590 nm, respectively. Mitochondrial damage is indicated by a decrease in the red/green fluorescence intensity ratio.

### Intracellular Reactive Oxygen Species (ROS) assays

The levels of intracellular ROS were measured using ROS probe dichlorodihydrofluorescein diacetate (DCFH-DA; Beyotime Biotech, Shanghai, China) and nuclear dye Hoechst 33342. Liver organoids were dissociated into single organoids and incubated with HCM containing 10 μM DCFH-DA and 10 μM Hoechst 33342 for 30 minutes at 37 °C in the dark. After incubation, the organoids were washed three times and resuspended in PBS with Antifade Mounting Medium before being observed using inverted fluorescence microscopy. DCFH-DA was visualized with the green fluorescence channel and Hoechst 33342 was visualized with the blue fluorescence channel. The organoids suspension was then transferred to a 96-well or 384-well microtiter plate, and the fluorescence intensity was detected on SpectraMax iD3 microplate reader. The excitation wavelength and emission wavelength of Hoechst 33342 are 350 nm and 460 nm, respectively. The excitation and emission wavelengths of DCFH-DA are 488 nm and 525nm, respectively. ROS level is indicated by changes in the green/blue fluorescence intensity ratio.

### Cell viability Assays

Cell viability of liver organoids was assessed using Calcein/PI Cell Viability/Cytotoxicity Assay Kit (Beyotime Biotech, Shanghai, China) following the manufacturer’s instructions. Liver organoids were dissociated into single organoids and incubated with Calcein AM/PI work solution for 30 minutes at 37 °C in the dark. Following incubation, Antifade Mounting Medium was added, and the organoids were observed by inverted fluorescence microscopy. Calcein AM was visualized using the green fluorescence channel, while PI was visualized using the red fluorescence channel. The organoids suspension was then transferred to a 96-well or 384-well microtiter plate, and the fluorescence intensity was detected on SpectraMax iD3 microplate reader. The excitation wavelength and emission wavelength of Calcein AM are 494 nm and 530 nm, respectively, while the excitation and emission wavelengths of PI are 535 nm and 617 nm, respectively. Changes in the green/red fluorescence intensity ratio indicate the cell viability of liver organoids.

### Statistical Analysis

The data were analyzed using GraphPad Prism 8.0 statistical software and expressed as mean ± Standard Error of Mean (SEM). Student’s t-test was used to evaluate the significance between the two groups. A one-way analysis of variance (ANOVA) was applied to compare the differences among three or more groups, with the Dunnett test used for comparison between multiple experimental and control groups. The *p-value* below 0.05 was considered statistically significant.

## Results

### Generation of liver organoids from hiPSCs

In this study, we established a differentiation protocol to transform hiPSCs into liver organoids by mimicking embryonic liver development. The protocol entailed the strategic addition of cytokines and small molecule compounds to the medium combined with Matrigel as a scaffold in a 3D culture (Figure 1A). Initially, the hiPSCs were treated with Activin A and BMP4 for 3 days under 2D culture conditions to drive their differentiation into Definitive Endoderm (DE) cells [15, 16]. Subsequently, the DE cells were treated with CHIR99021 and FGF-10 for 3 days to differentiate into foregut endoderm (FG) cells [17, 18]. FG cells were then isolated into single cells and implanted into Matrigel and 3D cultured for 6-7 days [17, 19]. During this period, spherical cell clusters composed of hepatic progenitor cells and irregular cell clusters composed of mesenchymal cells gradually appeared in Matrigel. Finally, cells in Matrigel were cultured in hepatocyte medium (HCM) supplemented with Dexamethasone, Oncostatin M, and HGF, leading to the gradual formation of liver organoids containing multiple liver cell types and 3D tissue architecture as early as day 22 (Figure 1A). Immunofluorescence (IF) and Western blotting (WB) were used to detect the expression of marker proteins at different stages of differentiation, further confirming the success of liver organoid differentiation (Figure 1B, C, D).

**Figure 1.**
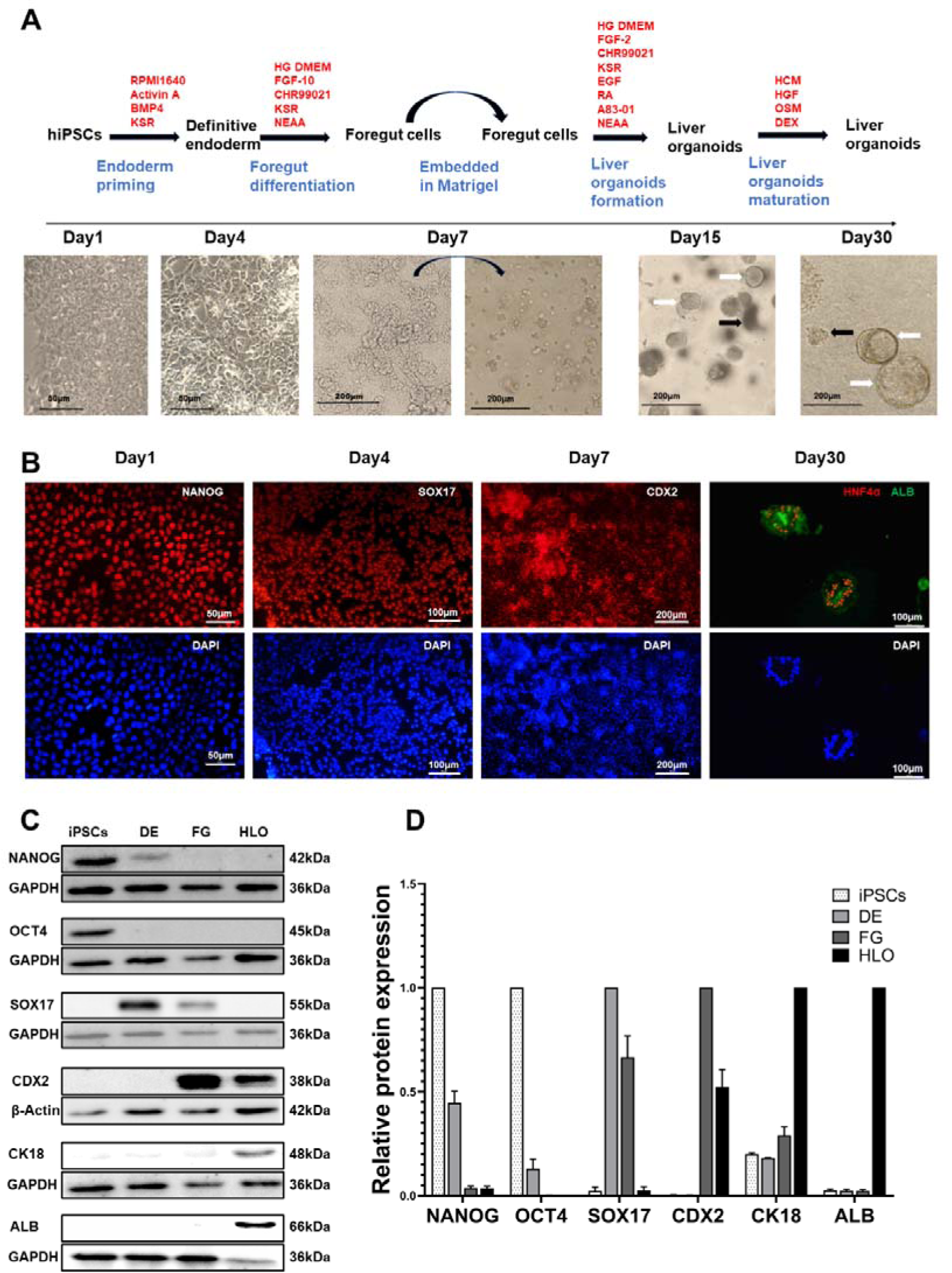
Differentiation of hiPSCs into liver organoids. (A) Schematic diagram of the differentiation protocol of liver organoids and morphological diagrams of different stages of differentiation. The white arrows showed a spherical mass, and the black arrows showed irregular mass in the bright field image on day 15 and day 30. (B) The immunofluorescence staining images showed the expression of pluripotent stem cell marker proteins NANOG, the definitive endoderm cells specific protein SOX17, the foregut cells specific protein CDX2, and hepatocytes specific proteins HNF4α and ALB at different stages of liver organoid differentiation. (C) Western blot also showed that the pluripotent stem cells marker proteins OCT4 and NANOG, the definitive endoderm cells specific protein SOX17, the foregut cells specific protein CDX2, and hepatocytes specific proteins CK18 and ALB were expressed at different stages of liver organoid differentiation. (D) The protein levels displayed in the histograms were determined by normalizing to the internal control products GAPDH or β-Actin. Data are shown as the mean ± SEM (n=3). The scale bar is shown in the picture. DE, Definitive endoderm; FG, Foregut; iPSCs, human induced Pluripotent Stem Cells; HLO: human liver organoids; ALB: Albumin; CK18: Cytokeratin 18; HNF4α, Hepatocyte nuclear factor 4 alpha.

### Identification of morphological structure and cellular components of liver organoids

Under the microscope, two types of structures are observed in the established liver organoids: spherical cell clusters and irregular cell clusters (Figure 1A). However, in paraffin sections, irregular cell clusters are dispersed around spherical cell clusters due to their disrupted loose structure. Immunofluorescence staining was further conducted on these cell clusters to determine their structure and cellular components. Firstly, paraffin sections of liver organoids were immunofluorescent stained for the hepatocyte marker hepatocyte nuclear factor (HNF4α), epithelial cell markers E-cadherin and cytoplasmic junction protein (zonula occludens-1, ZO-1). HNF4-α, a transcription factor that regulates a variety of liver genes, is located in the nucleus of hepatocytes and serves as a marker of hepatocyte lineages. E-cadherin, localized on the cell surface and at cell-to-cell junctions, is a marker of epithelial cells in the liver. ZO-1, a tight junction mucin, is mainly situated in the lumen wall formed by epithelial cells. The staining results showed that the spherical cell clusters in the liver organoids were hollow saccular organoids composed of epithelial cells (hepatocytes and cholangiocytes) with polar epithelial structures (Figure 2A). Furthermore, immunofluorescence staining with the cholangiocyte marker Cytokeratin19 (CK19) demonstrated that some of the organoids with the spherical cystic structure contained hepatocyte-like-cells, some contained cholangiocyte like cells, and some were composed of liver progenitor cells with bipotential of hepatocyte and cholangiocyte (Figure 2A).

**Figure 2.**
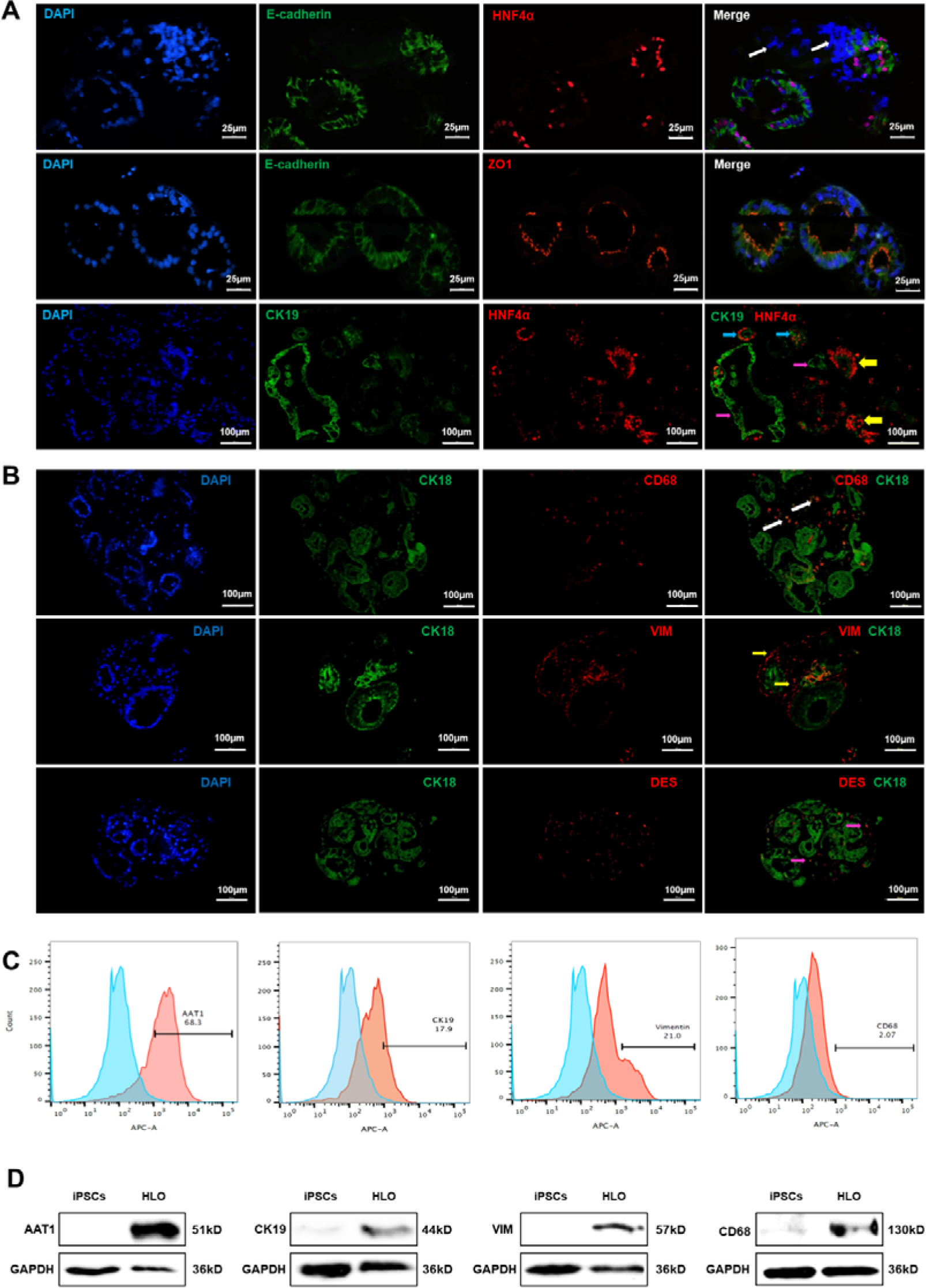
Identification of liver organoid’s structure and cell composition. (A) Paraffin section immunofluorescence staining showed that the spherical mass was composed of epithelial cells, and the irregular mass was non-epithelial mesenchymal cells. White arrows showed mesenchymal cells, yellow arrows showed hepatocytes, purple arrows showed cholangiocytes, and blue arrows showed bipotent hepatic progenitor cells. (B) Paraffin section immunofluorescence staining showed that stromal cells contained the Kupffer cell marker CD68 and hepatic stellate cell-specific proteins VIM, DES, as indicated by white arrows, yellow arrows, and purple arrows, respectively. (C) Flow cytometry confirmed the presence of CD68^+^, CK19^+^, AAT1^+^, and VIM^+^ cell populations in liver organoids. (D) Western blot showed that the hepatocytes specific protein AAT1, the cholangiocytes specific protein CK19, the Kupffer cell marker protein CD68, and hepatic stellate cell-specific protein VIM were expressed in liver organoids. The scale bar is shown in the picture. iPSCs, human induced Pluripotent Stem Cells; HLO: human liver organoids; AAT1: α1-antitrypsin; ZO-1: zonula occludens-1; VIM: vimentin; DES: desmin; CK18: Cytokeratin 18; CK19: Cytokeratin 19; HNF4α, Hepatocyte nuclear factor 4 alpha.

Immunofluorescence staining revealed that the irregular cell clusters scattered around the spherical structure were negative for E-cadherin, ZO-1, and HNF4-α, indicating their non-epithelial mesenchymal cell nature. Further immunofluorescence staining showed that these irregular stromal cell clusters expressed hepatic stellate cell-specific protein or markers, including Vimentin (VIM), and Desmin (DES), as well as the Kupffer cells marker CD68 (Figure 2B). Flow cytometry also detected the presence of liver-specific proteins such as α1-antitrypsin (AAT1) positive cells, CK19 positive cells, VIM positive cells, and CD68 positive cells in the organoids (Figure 2C). Western blotting further confirmed the expression of HNF4-α, CK19, Vimentin, and CD68 in liver organoids (Figure 2D). These findings indicate that, in addition to hepatic progenitor cells capable of differentiating into hepatocytes and cholangiocytes, the liver organoids we established also contain hepatic stellate cells and Kupffer cells.

### Assessment of liver-specific protein expression and characterization of liver-specific function in the liver organoids

We subsequently performed immunofluorescence staining and Western blotting to assess the expression of liver-specific proteins in the liver organoids. The results demonstrated the presence of liver-specific proteins including Albumin (ALB), α1-antitrypsin (AAT1), Cytokeratin 18 (CK18), alcohol dehydrogenase (ADH), alkaline phosphatase (ALP), and cytochrome P450 family member 2E1 (CYP2E1) in the liver organoids (Figure 3A, B).

**Figure 3.**
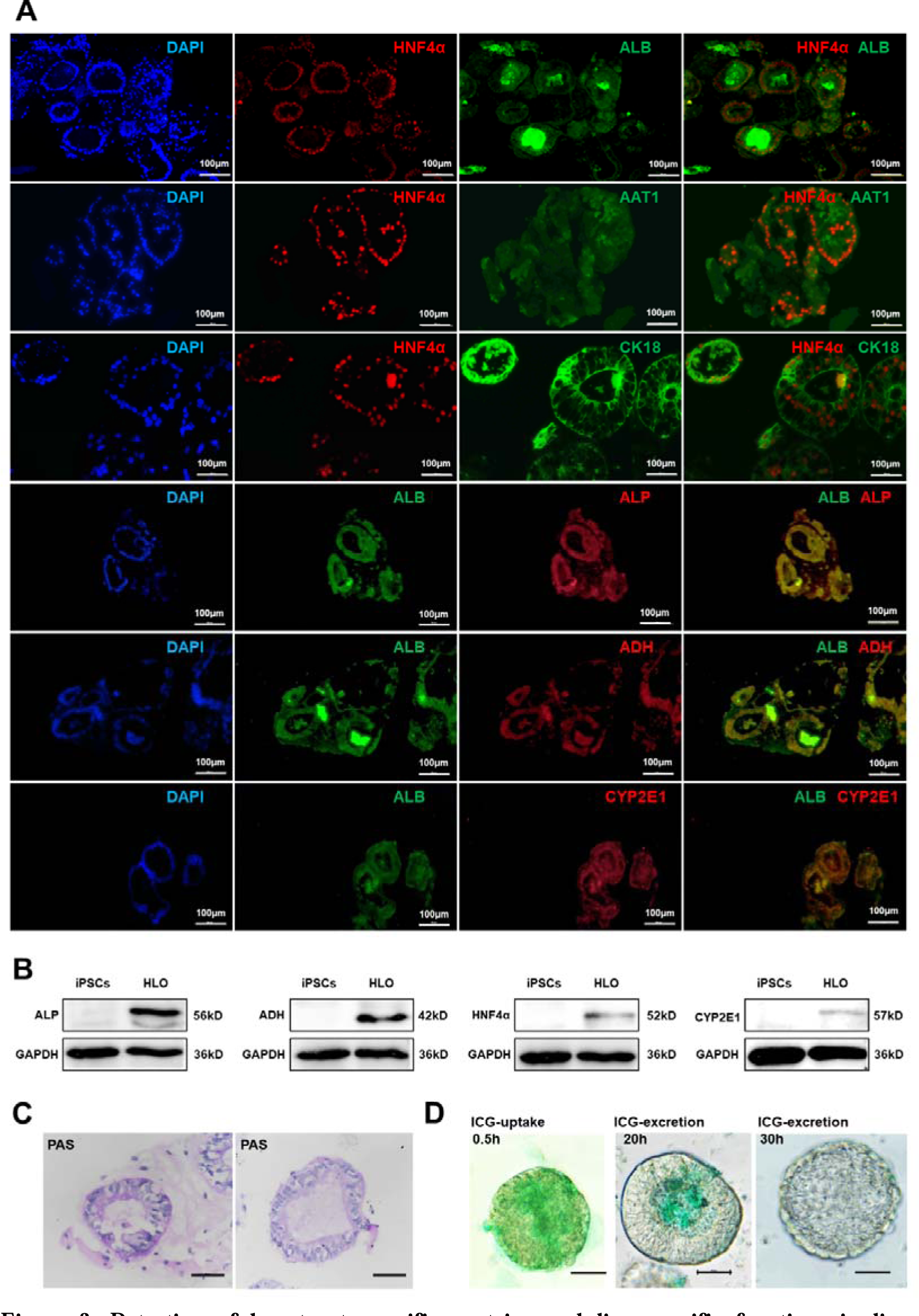
Detection of hepatocyte-specific proteins and liver-specific functions in liver organoids. (A) Paraffin section immunofluorescence staining showed the expression of hepatocyte-specific proteins HNF4α, ALB, AAT1, CK18, ADH, ALP, and CYP2E1 in liver organoids. The scale bar is 100 μm. (B) Western blot also confirmed the expression of hepatocyte-specific proteins ALP, ADH, HNF4α, and CYP2E1 in liver organoids. (C) Paraffin section PAS staining confirmed that liver organoids could synthesize and store glycogen. The scale bar is 50 μm. (D) ICG experiments confirmed that liver organoids can uptake and excrete ICG. The scale bar is 50 μm. iPSCs, human induced Pluripotent Stem Cells; HLO: human liver organoids; ALB: Albumin; AAT1: α1-antitrypsin; CK18: Cytokeratin 18; ADH: Alcohol dehydrogenase; ALP: Alkaline phosphatase; HNF4α, Hepatocyte nuclear factor 4 alpha; CYP2E1: cytochrome P450 family member 2E1; PAS: Periodic Acid-Schiff staining; ICG: Indocyanine Green.

Subsequently, we performed periodic acid Schiff stain (PAS) and Indocyanine green (ICG) uptake and release assay on the liver organoids. PAS staining is mainly used to detect sugars in tissues and can indicate glycogen storage in liver organoid cells. PAS staining of liver organoids paraffin sections showed positive staining in hepatocytes, indicating the liver organoids’ capability to synthesize glycogen (Figure 3C). The ICG uptake and release assay assesses the functionality of hepatocytes by tracking the binding of ICG with albumin, its uptake by hepatocytes, and subsequent excretion. ICG experiments showed that ICG was rapidly taken up by hepatocytes in the liver organoids within half an hour of treatment. Upon switching to a normal HCM medium, ICG was gradually excreted from the cells and was completely removed after 30 hours. This confirmed the presence of functional hepatocytes in the liver organoids with both ICG uptake and excretion capabilities (Figure 3D).

### Establishment of ALD model in liver organoids and analysis of associated gene expression

The ALD model was established by introducing different concentrations of ethanol into the culture medium between days 27-30 of liver organoid differentiation, while a control group (without ethanol) was maintained concurrently. Liver organoids were harvested after 72 hours of ethanol treatment, and RNA was extracted to obtain the liver organoid cDNA library through reverse transcription PCR. The expression of liver injury-related gene in the ALD model was assessed via fluorescence quantitative PCR and compared with the control group (Figure 4). The results showed that upon treatment with ethanol at 100 mM and 200 mM, the expression of genes *ACC1, FASN,* and *SCD* associated with lipid metabolism increased in liver organoids. However, their expression remained unchanged after treatment with 400 mM ethanol (Figure 4A). Similarly, the expression of genes *COL1A1, COL3A1, ACTA2, DES, TGF*β*1,* and *VIM* involved in liver fibrosis increased in liver organoids treated with 100 mM and 200 mM ethanol. Conversely, their expression levels remained unchanged, and in some cases decreased, following treatment with 400 mM ethanol (Figure 4B). Furthermore, the expression of genes *ADH1* and *CYP2E1,* which encode enzymes involved in alcohol metabolism, increased in liver organoids post-ethanol treatment. However, the expression of genes encoding cytochrome oxidase *CYP2C9* and *CYP3A4* remained unchanged (Figure 4C). In addition, the expression levels of interleukin encoding genes *IL-10, IL-3, IL-6, IL-1*β, and *IL-17* in liver organoid cells showed varying degrees of increase following ethanol treatment (Figure 4D). Nonetheless, due to potential variability between different experimental batches, some changes in gene expression did not reach statistical significance.

**Figure 4.**
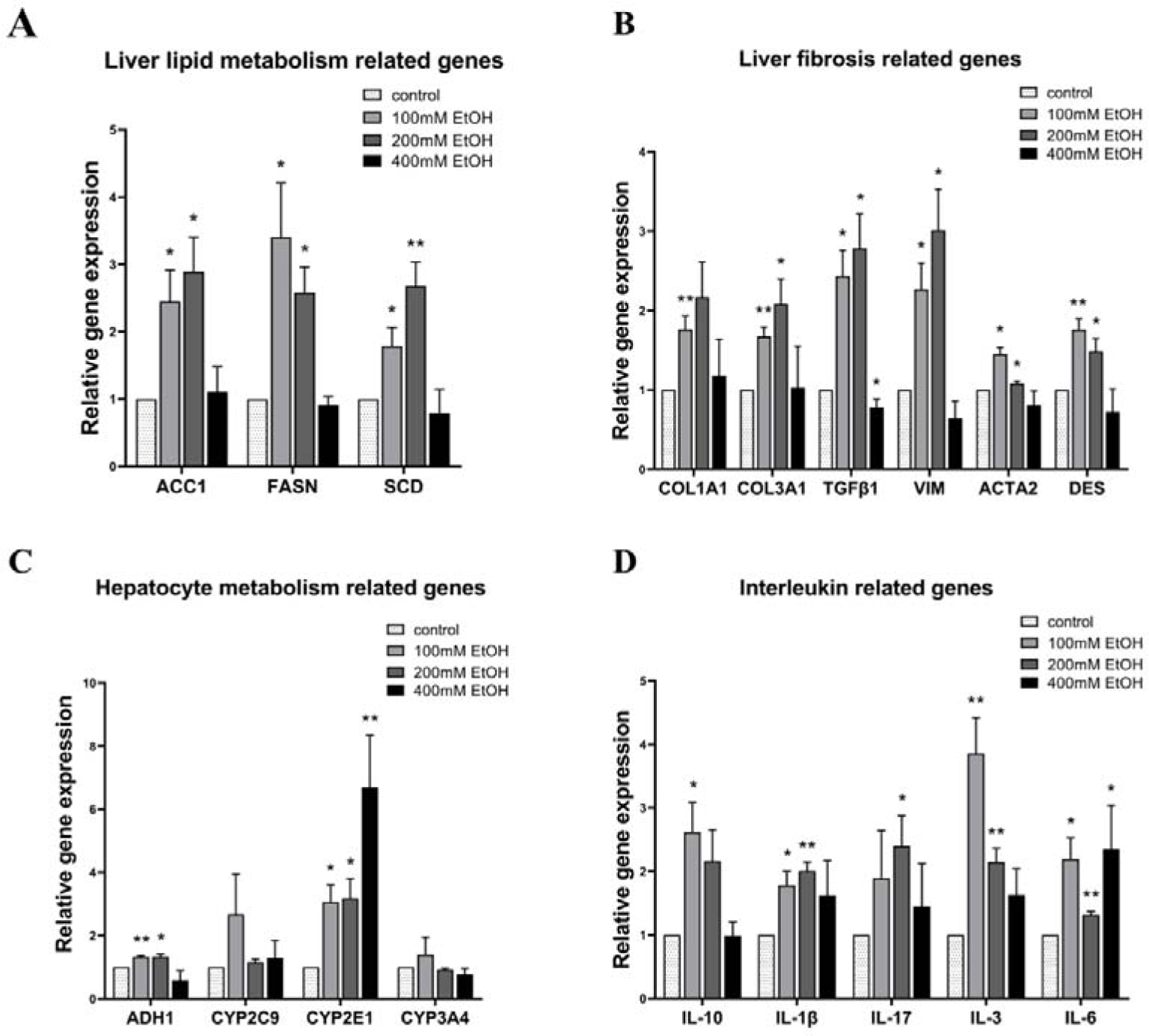
Gene expression in liver organoids after treatment with different ethanol concentrations. The expression levels of liver lipid metabolism-related genes (A), liver fibrosis-related genes (B), hepatocyte metabolism-related genes (C), and interleukin-related genes (D) in liver organoids treated with different concentrations of ethanol were detected by qPCR, and compared with the expression levels in liver organoids of the control group. *GAPDH* was used as an internal reference. Data are from 3 samples of 3 independent differentiation experiments and are shown as mean ± SEM. Analysis of variance was used to detect whether there was a statistical difference in the genes’ expression levels among the groups. When the difference was statistically significant, *: *p*<0.05; **: *p*<0.01. Control: Liver organoids in the control group; EtOH: Liver organoids in different concentration ethanol treated groups.

### Pathological changes in the ALD model

The liver organoids were subjected to ethanol treatment to establish an ALD model. After 24 hours of treatment, the mitochondrial membrane potential (MMP) and levels of reactive oxygen species (ROS) in the model were detected using a multi-function microplate reader and compared with the control group of liver organoids. The results showed that following 24 hours of ethanol treatment, the MMP decreased and ROS levels increased in organoid cells (Figure 5A, B). Consistently, under the fluorescence microscope, both JC green fluorescence intensity and DCF green fluorescence intensity in liver organoids increased after ethanol treatment, further confirming the decrease in MMP and increase in ROS (Figure 5A, B). After 48 hours of ethanol treatment, the levels of neutral lipids in the model were detected with BODIPY staining and compared with the control group liver organoids. Observation under the fluorescence microscope revealed an increase in BODIPY green fluorescence intensity in the liver organoids after ethanol treatment, indicating that the level of neutral lipids in the organoid cells increased (Figure 5C). The results of the microplate reader also showed that after 48 hours of ethanol treatment, the ratio of green/blue fluorescence intensity in liver organoids increased, indicating that the level of neutral lipids in organoid cells increased (Figure 5C). After 72 hours of ethanol treatment, the activity and death of cells in the organoids were detected by Calcein/PI Cell Viability and Cytotoxicity Detection Kit. Under the fluorescence microscope, the red fluorescence intensity increased in the liver organoids after ethanol treatment, indicating that the dead cells increased in the organoid (Figure 5D). Similarly, the results from the multi-microplate reader showed that the ratio of red/green fluorescence intensity in liver organoids increased after 72 hours of ethanol treatment, indicating a decrease in organoid cell activity and an increase in the proportion of dead cells (Figure 5D). Nonetheless, some changes may not have reached statistical significance, possibly due to significant errors between different experimental batches.

**Figure 5.**
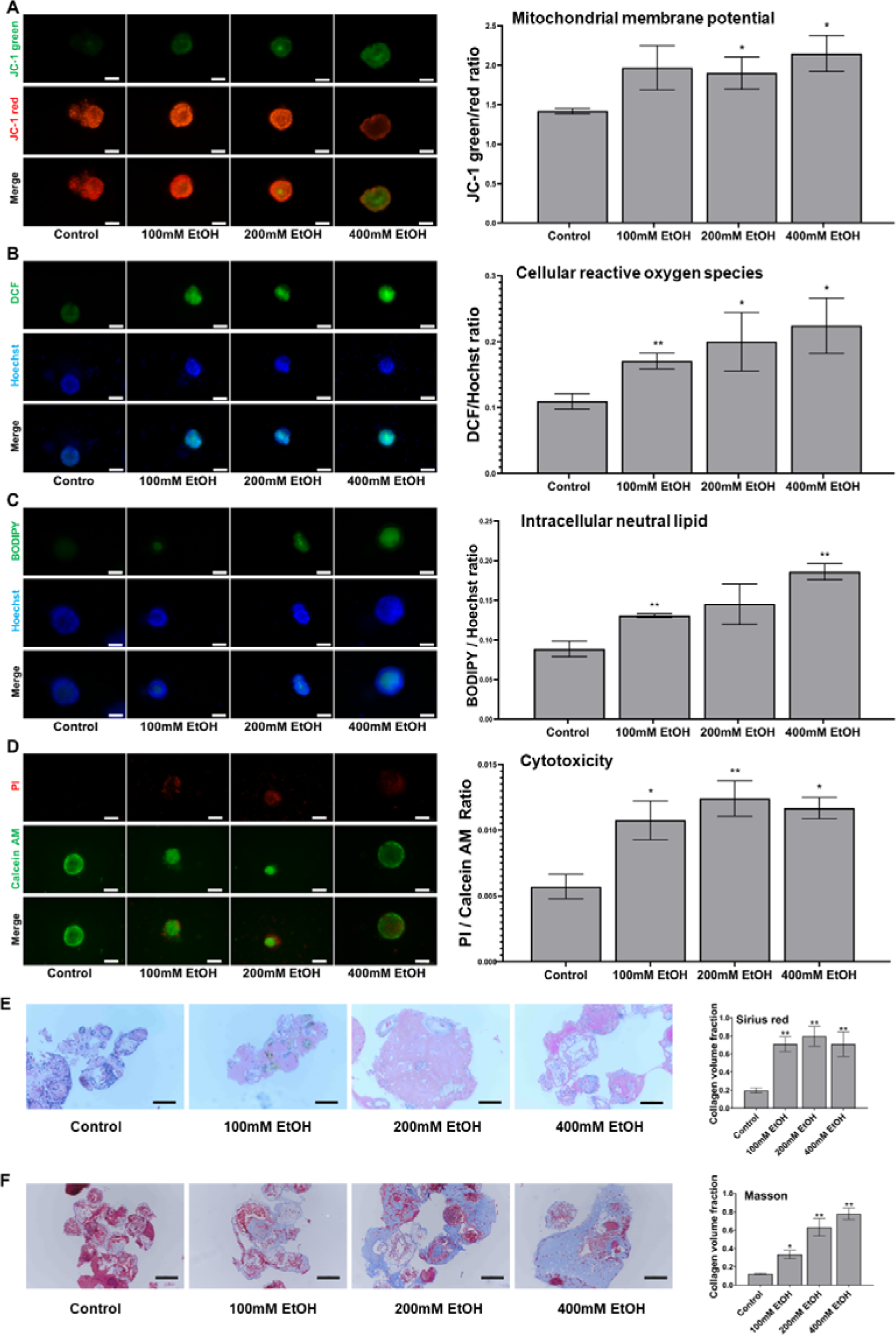
Pathological damage in ALD model of liver organoids. (A-D) The MMP (A) and cellular ROS (B) levels in liver organoids after ethanol treatment for 24 hours, neutral lipid levels (C) after ethanol treatment for 48 hours, and the dead cells proportion (D) after ethanol treatment for 72 hours were detected and compared with the results of liver organoids in the control group. Left: Images of HLO under the fluorescence microscope. Scale bar: 100 μm. Right: Graphs showed the fluorescence intensity ratios detected by SpectraMax iD3 microplate reader. Data are presented as mean ± SEM (n = 3 per group). Analysis of variance was used to detect whether there was a statistical difference between each ethanol-treated group and the control group. *: *p*<0.05; **: *p*<0.01. (E, F) The paraffin sections of liver organoids from the control and ethanol-treated groups were stained with Sirius red (E) and Masson staining (F). The collagen volume fractions are shown as mean ± SEM (n = 3 per group). Analysis of variance was used to detect whether there was a statistical difference between each ethanol-treated group and the control group. *: *p*<0.05; **: *p*<0.01. Scale bar: 100 μm. Control: Liver organoids in the control group; EtOH: Liver organoids in different concentration ethanol treated groups.

Furthermore, after 72 hours of ethanol treatment, liver organoids from each group were harvested to prepare paraffin sections for Sirius red staining and Masson staining. The results showed an increased collagen volume fraction in liver organoids following ethanol treatment (Figure 5E, F).

### The therapeutic impact of drugs on the liver organoids model of ALD

Finally, we assessed this model with three drugs reported in the literature to have therapeutic effects on ALD to verify its suitability for drug screening. On the 27th to 30th day of liver organoid differentiation, 100 mM ethanol was added to the culture medium to establish the ALD model. Meanwhile, low, medium, and high three doses of interleukin-22 (IL-22), Metadoxine (Met) and N-acetylcysteine (NAC) were added to the medium, respectively, to treat the ALD model. After three days, the therapeutic effect of the drug on the ALD model was assessed using the Calcein/PI cell viability and cytotoxicity detection kit with a multifunctional microplate reader. Simultaneously, the toxicity of the drug to the liver organoids was also evaluated.

The results indicated that the cell viability of liver organoids after treatment with three drugs did not significantly differ from that in the control group of liver organoids without drug treatment, suggesting that these drugs at their respective doses had no toxicity to liver organoids. After treatment with 100 mM ethanol, the proportion of dead cells in the liver organoids increased and the cell viability decreased significantly, confirming the successful establishment of the ALD model (Figure 6A, B, C). The cell viability of the IL-22 treatment group was higher than that of the untreated group, and the cell viability of the high-dose group was not different from that of the control group, indicating that IL-22 exerted preventive and therapeutic effect on ALD, with a more pronounced effect at high-doses (Figure 6A). Similarly, the cell viability of the Met treatment group was also higher than that of the untreated group. The middle and high dose groups showed differing cell viability compared to the control group, indicating that Met had preventive and therapeutic effects on ALD, with the medium treatment doses showing the most significant therapeutic effect (Figure 6B). In the NAC treatment group, cell activity was higher than in the untreated group, with the medium dose group showing no difference from the control group. This suggests that NAC had both preventive and therapeutic effects on ALD, with the medium dose being the most effective (Figure 6C).

**Figure 6.**
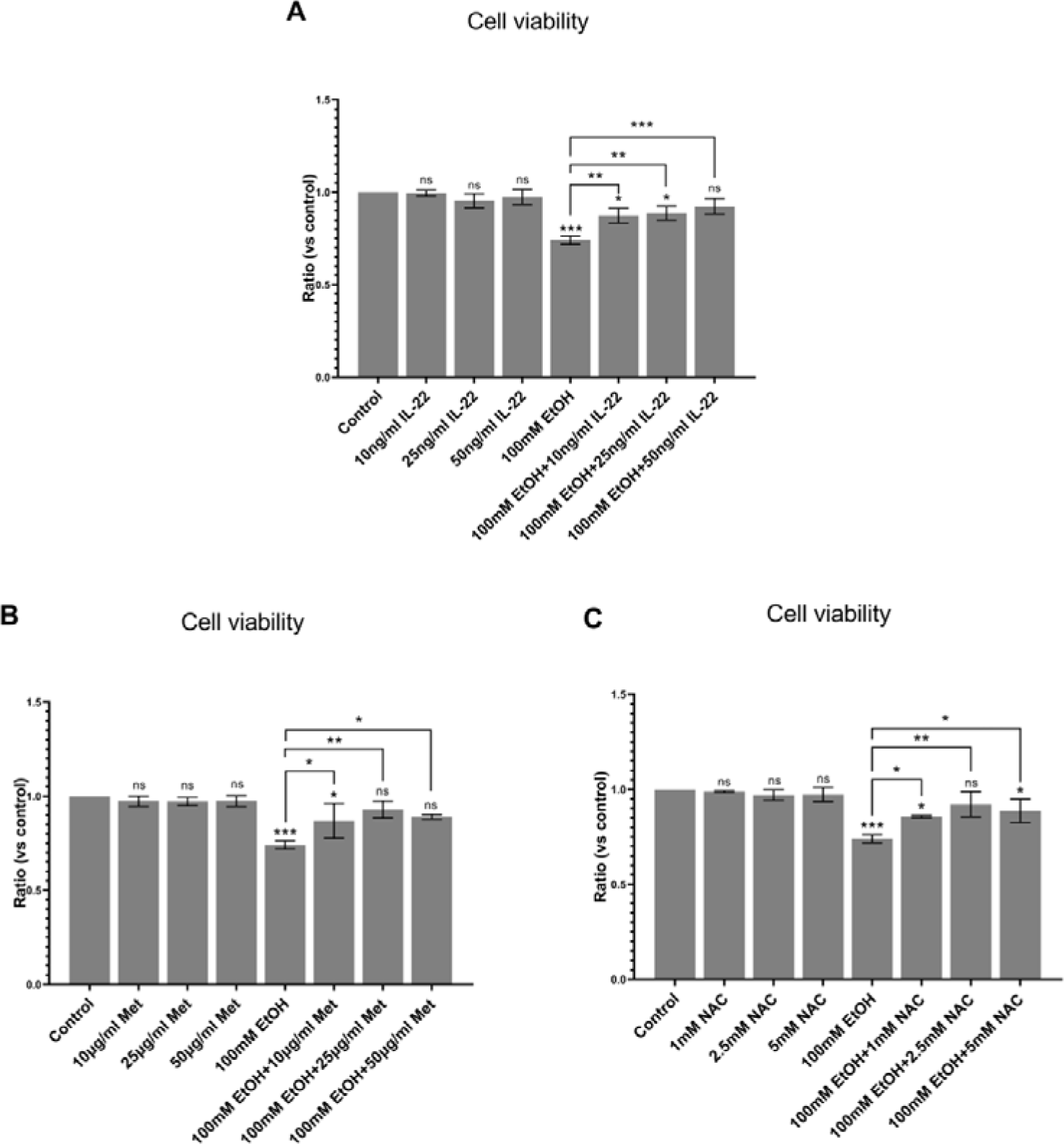
Drug treatment of ALD models. The ALD liver organoid model was established with 100 mM ethanol treatment and the model was treated with different doses of IL-22, Met, and NAC, respectively. Three days later, the therapeutic effect was detected with Calcein/PI Cell Viability and Cytotoxicity Detection Kit, and the toxicity of the three drugs to liver organoids was also detected. (A) The ALD model was treated with IL-22 at doses of 10 ng/ml, 25 ng/ml, and 50 ng/ml, respectively. (B) The ALD model was treated with Met at doses of 10 μg/ml, 25 μg/ml, and 50 μg/ml, respectively. (C) The ALD model was treated with NAC at 1 mM, 2.5 mM, and 5 mM, respectively. Data are from 3 independent samples and are shown as mean ± SEM. Analysis of variance was used to detect whether there was a statistical difference between each medicine-treated group and the control group or medicine-untreated ALD group. ns: no statistical significance; when the difference was statistically significant, *: *p*<0.05; **: *p*<0.01, ***: *p*<0.001. IL-22: interleukin 22; Met: Metadoxine; NAC: N-acetylcysteine; Control: Liver organoids in the control group; EtOH: Liver organoids in different concentration ethanol treated groups.

## Discussion

The liver, the largest solid organ in the human body, plays a crucial role in metabolism [20]. Liver diseases have emerged as a prominent cause of mortality worldwide [21]. Among these, alcoholic liver disease (ALD) stands as a common liver disease [1]. Nevertheless, the advancement of ALD research is impeded by the lack of suitable disease models. In laboratories, researchers often establish ALD models of rats or mice through alcohol intragastric administration, feeding alcoholic diet, or intraperitoneal injection of alcohol [5, 6]. While these models can replicate some pathological manifestations of human alcoholic liver injury, such as hepatocyte steatosis, liver necrosis, apoptosis and inflammation, species-specific variations persist [5, 6]. The metabolic pathways of alcohol in humans and model animals are not entirely the same, as do their response to liver injury [8]. Therefore, findings from animal models may not fully reflect the complexities of human physiology, limiting their utility in ALD research [8].

In addition, human liver cell lines, human liver cancer cell lines, and primary human hepatocytes (PHHs) have also been employed to establish ALD models. Typically, these models involve the addition of ethanol to the cell culture medium to mimic alcohol’s impact on liver cells *in vivo* [22–24]. However, these 2D cultured cells exhibit limitations. They consist of a single component, lack intricate cell interactions, and fail to replicate the native characteristics of the liver microenvironment. Furthermore, once PHHs are removed from their *in vivo* microenvironment, they rapidly degenerate, and lose their original functionality, and experience diminished proliferative capacity [25]. The liver cell lines are difficult to culture, and recent reports have raised concerns about contamination with HeLa cell lines [26], questioning the reliability of experimental results. The liver cancer cell lines are abnormal cells with functional disparities compared to normal liver cells, necessitating cautious interpretation of results. Consequently, these 2D cultured cell models face challenges in reproducing the pathological manifestations of ALD *in vivo*. These limitations, combined with individual differences in susceptibility to human ALD, limit the utility of these cell models[9, 10].

Moreover, the intricate interplay among hepatocytes, immune cells, and stellate cells, critical in the pathogenesis of ALD [1, 27], presents a challenge for the *in vitro* culture model composed of these individual cell components. Such models struggle to faithfully replicate the full spectrum of pathological changes observed in alcoholic liver injury *in vivo*. While some researchers have co-cultured hepatocytes with liver non-parenchymal cells to simulate liver function and microenvironment [28, 29], this approach introduces complexity to cell culture. In a previous study, researchers co-cultured human liver organoids differentiated from human embryonic stem cells (ESCs) with human fetal liver mesenchymal cells (hFLMCs) in 3D culture and established an ALD model by ethanol treatment [30]. This ALD model contains a variety of liver cell components and effectively recapitulates certain pathological manifestations of ALD [30]. However, it is important to note that the study is technically complicated and time-consuming, and hFLMCs are not easy to obtain, significantly limiting their application.

In this study, we employed a sophisticated approach involving the introduction of cytokines and small molecule compounds to the cell culture medium, combined with 3D culture. This method allowed us to successfully differentiate hiPSCs into liver organoids in just about 22 days. These liver organoids exhibited a comprehensive composition, containing hepatocyte-like cells, cholangiocyte-like cells, hepatic stellate cells, and Kupffer cells. Various measurements confirmed that the liver organoids possess liver cell polarity, multiple liver-specific functions, and express various liver-related proteins. Subsequently, we harnessed this advanced liver organoid platform to establish a human ALD model by treating liver organoids with ethanol. By fluorescence quantitative PCR, we found that genes related to ALD-related liver injury were upregulated in the ALD liver organoid model, such as alcohol metabolism genes, lipid metabolism genes, fibrosis-related genes, and inflammation-related interleukin genes. In hepatocytes, ethanol is mainly metabolized by alcohol dehydrogenase and CYP2E1 into acetaldehyde, which is responsible for the generation of ROS. ROS cause oxidative stress and steatosis, which lead to inflammation and apoptosis [1, 4, 31]. Meanwhile, ethanol can also induce liver cell apoptosis by activating the intrinsic mitochondrial apoptosis pathway. Furthermore, hepatocyte death, followed by a release of damage-associated molecular patterns (DAMPs), can activate Kupffer cells[1]. Activated Kupffer cells produce many proinflammatory cytokines (for example, interleukins) that promote HSCs activation. ROS and acetaldehyde can also directly stimulate HSCs, resulting in fibrogenesis[4]. Remarkably, our model recapitulated key pathological features of ALD *in vitro*, including hepatocyte steatosis, mitochondrial damage, increased ROS level, hepatocyte necrosis, and fibrosis. Finally, we assessed this model with three drugs reported in the literature to have therapeutic effects on ALD. NAC and Metadoxine are antioxidants, which can counteract the oxidative effects caused by ethanol [32, 33]. Interleukin-22 can induce proliferation genes and reduce apoptosis genes in hepatocytes [34]. The screening results identified the therapeutic effects of these three drugs on ALD, confirming that the model is suitable for large-scale drug screening.

Evidence suggests that susceptibility to ALD is influenced by genetic factors, and the pathological processes and drug responses are individual-specific [35]. To address this variability, the liver organoids derived from patient-specific hiPSCs are particularly valuable. These organoids carry the patient’s genetic information, providing a more accurate reflection of how the patient’s liver responds to alcohol and drugs. Additionally, since they are not rejected by the body’s immune system, the hiPSCs-derived liver organoids can be used as a potential source of cell therapy and for liver transplantation. Finally, liver organoids are suitable for large-scale and engineering culture [11, 13]. Therefore, liver organoids established from patient-derived hiPSCs are more suitable as a drug screening platform for ALD, which can be used to evaluate the efficacy and toxicity of candidate drugs and predict the possibility of severe side effects. Altogether, these advantages underscore the immense potential of hiPSCs-derived liver organoids in personalized drug development and liver transplantation therapy.

